# SPARP-seq reveals poly(A) tail shortening as a hallmark of the active human translatome

**DOI:** 10.1101/2025.11.25.690357

**Authors:** Yi Shu, Juzuo Li, Zhiyuan Sun, Yanping Li, Wenqin Lu, Yanping Long, Xi Chen, Sisi Li, Wei Chen, Jixian Zhai

## Abstract

The poly(A) tail at the 3’ end of mRNAs plays a critical role in the synthesis of proteins, but its dynamic relationship with the translation process has been unclear. Here, using a novel single-molecule approach SPARP-seq that integrates poly(A) assessment with ribosome profiling, we demonstrate that poly(A) tail shortening is a universal hallmark of active translation in human cells. We find that most mRNAs begin translation on single ribosome (monosome) with long tails that progressively shorten as additional ribosomes are recruited. In contrast, highly expressed housekeeping genes are translated efficiently on monosomes and possess intrinsically short poly(A) tails. Furthermore, we find that translationally repressed transcripts, such as those retaining introns, are trapped on monosomes with long tails. Pharmacological inhibition of translation induces global poly(A) tail lengthening, confirming that deadenylation is coupled to ongoing translation. These findings reveal a dynamic system where the poly(A) tail integrates mRNA status with translational fate, defining the monosome as a central hub for post-transcriptional control.

## Introduction

Translation is a fundamental step in gene expression, serving as the critical bridge between genetic information and functional protein synthesis. This process occurs on ribosomes, which exist either as mono-ribosome (monosome) or as poly-ribosome (polysome), where multiple ribosomes simultaneously engage the same mRNA template^1–3^. A key regulator of mRNA fate, the 3’ poly(A) tail is critical for enhancing translation initiation, mRNA stability and nuclear export^4–13^. The interaction between the poly(A)-binding protein (PABPC) and the 5’ cap-binding complex eIF4F circularizes the mRNA, promoting efficient ribosome recruitment and recycling^14–17^. Despite the established importance of the poly(A) tail, the clear correlation between poly(A) tail length and translation observed in early embryos is not maintained in somatic cells^4,8,18,19^. The paradoxical relationship between the poly(A) tail’s importance and its poor correlation with translation in somatic cells may be due to distinct patterns being averaged out by bulk measurements that lack single-molecule resolution of both the mRNA isoforms and its specific poly(A) tail lengths.

This complexity is compounded by an evolving understanding of ribosome function itself. For decades, polysome has been widely regarded as the primary sites of active protein synthesis, reflecting robust translational engagement. In contrast, monosome have been viewed as translationally inactive, often considered newly assembled ribosome or those dissociated from mRNA after translation termination^17,20–22^. However, this view has been challenged by ribosome profiling studies, which revealed that a significant fraction of monosome is actively engaged in elongation, particularly for certain classes of mRNAs such as highly expressed housekeeping genes in yeast and synaptic mRNAs in neuronal processes^23,24^. Despite this, the full spectrum of monosome function and, critically, how they might sort different classes of mRNA for distinct translational fates, remains unresolved.

Resolving these questions requires methods that can simultaneously determine an mRNA’s isoform structure, its poly(A) tail length, and its precise association with the translational machinery. Classic Ribosome profiling (Ribo-seq) has revolutionized the study of translation by mapping ribosome positions at codon resolution, providing genome-wide insights into translation efficiency, ribosome stalling, and new open reading frames^1–3,25,26^. However, its reliance on short-read sequencing prevents analysis of full-length isoforms or poly(A) tails. Recent advances in long-read sequencing technologies offer the potential to overcome this limitation. For instance, Frac-seq^27^, Ribo-STAMP^28^ integrate long reads with subcellular fractionation to measure alternative splicing during translation with isoform resolution. However, Ribo-STAMP’s enzymatic tagging requires the expression of a fusion protein, precluding its use in many primary tissues or clinical samples^28^. Direct RNA sequencing of fractionated RNA can measure RNA modification and tail lengths^29,30^, but suffers from high input requirements (∼500 ng of poly(A)+ RNA per library) and biases inherent in oligo-dT-based library preparation^31^. While specialized methods like FLAM-seq^32^, PAIso-seq^33^, and Nano3P-seq^34^ have been developed to quantify poly(A) tail features using long reads, they do not simultaneously resolve the translational state of the mRNA.

To overcome these specific and complementary limitations, we developed SPARP-seq (Single-molecule Poly(A)-informed Ribosome Profiling), a low-input, high-throughput, unbiased approach which is capable of simultaneously assessing full-length mRNA transcripts along with their poly(A) tail status across distinct ribosome populations. SPARP-seq library construction can begin with as little as 1 ng rRNA-depleted RNA (∼100 ng Ribosome-associated RNA) and maintains high reproducibility among biological replicates (Fig. 1a), making it widely applicable to various of samples with limited RNA quantities. By avoiding oligo-dT enrichment and genetic modification, SPARP-seq provides a high-resolution view of the active human translatome, enabling a direct investigation into the dynamic interplay between poly(A) tail length and translational control.

**Fig. 1.**
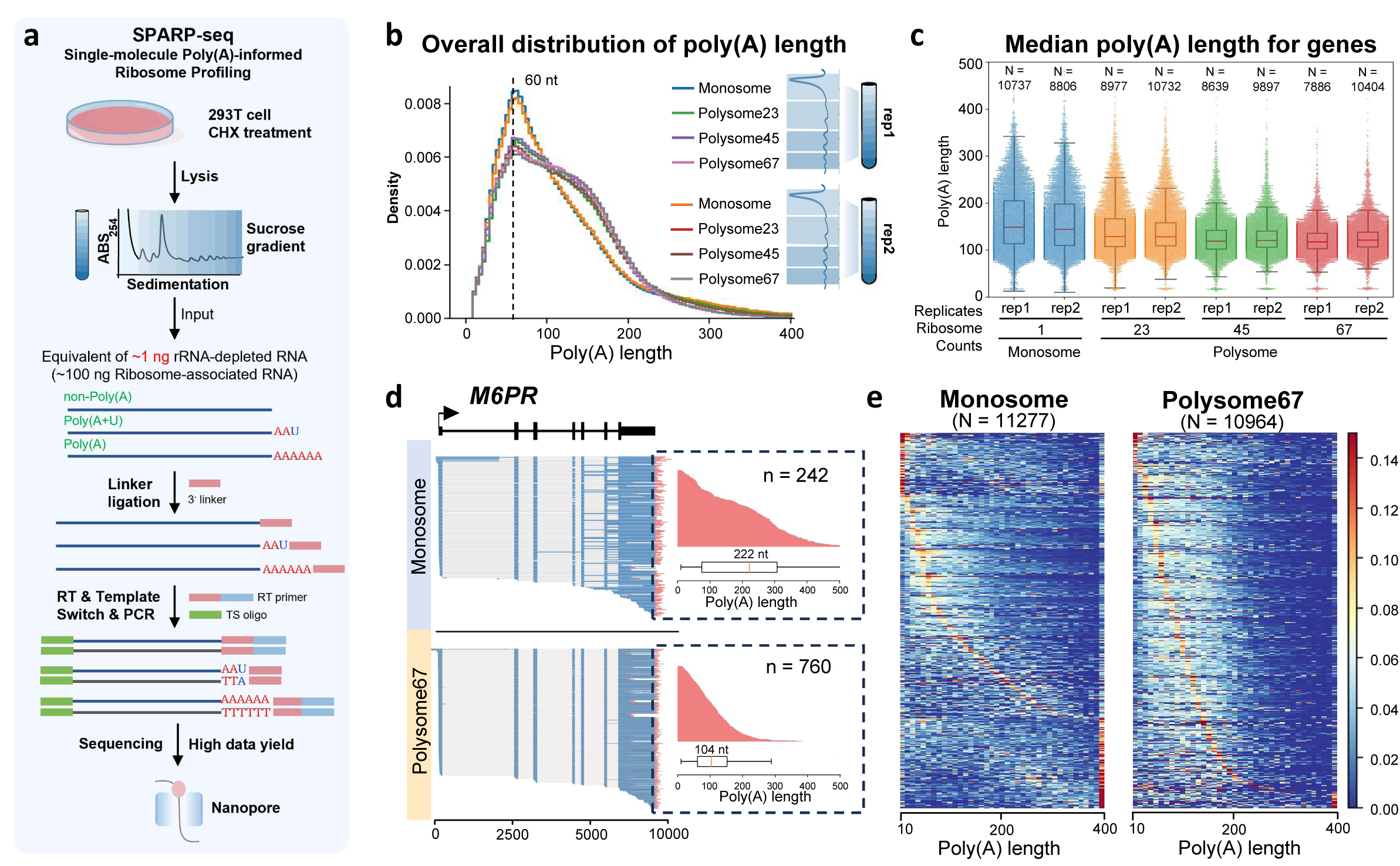
SPARP-seq captures the dynamics of poly(A) tail during translation. **a.** Schematic overview of the Single-molecule Poly(A)-informed Ribosome Profiling (SPARP-seq) workflow. 293T cells are treated with cycloheximide (CHX), lysed, and subjected to sucrose gradient centrifugation to separate mRNAs based on the number of bound ribosome. Ribosome-associated RNA is isolated, with an input equivalent of ∼1 ng of rRNA-depleted RNA, and a 3’ linker is ligated to the RNA molecules. The extracted RNA undergoes reverse transcription with a custom primer, template switching, and PCR, generating cDNA libraries for high-yield Nanopore sequencing. **b.** Overall poly(A) tail length distribution for all reads across different translational fractions (Monosome, Polysome23, Polysome45, Polysome67). Two replicates were included and only reads with poly(A) tails longer than 10 nt were selected for analysis. **c.** Boxplots showing the median poly(A) tail length at the gene level for each translational fraction from two biological replicates. Only genes with more than 10 uniquely mapped poly(A) reads were included in the analysis. **d.** Examples of single-molecule reads from the *M6PR* gene, showing read alignments and poly(A) tail length distributions in monosome and polysome67 fractions. **e.** Heatmap illustrating the poly(A) tail length patterns of all genes in monosome (N = 11,277) and polysome67 (N = 10,964) fractions. Data from two biological replicates within each fraction were merged. Each row in the plot represents the poly(A) tail length distribution of a single gene (10 nt bin).

## Results

### High-Resolution Profiling of Ribosome-Associated Transcripts by SPARP-seq

To resolve the paradoxical relationship between poly(A) tail length and translation, we developed SPARP-seq to simultaneously capture full-length transcript structure, their poly(A) tail features, and their association with distinct ribosome populations at single-molecule resolution. We fractionated lysates from human 293T cells by sucrose gradient centrifugation to separate into monosome and polysome fractions by ribosome count (1, 2–3, 4–5, or 6–7). RNA from each fraction was then subjected to a ligation-based library preparation protocol, followed by long-read sequencing (Fig. 1a). Deep sequencing of 12 libraries from two biological replicates yielded approximately 110 million reads using Oxford Nanopore Technologies platform (Table S1). Comparison of gene-level read counts and poly(A) tail length between the two replicates revealed high concordance, validating the robustness of our experimental approach (Supplemental Fig. 1-2).

Subsequently, we validated our method against parallel Ribo-seq experiments, the gold standard for measuring translation (Table S2). Data from two replicates on monosome, polysome and total cellular lysate (global) displayed the hallmarks of high-quality ribosome profiling, including robust three-nucleotide periodicity within open reading frames (ORFs) (Supplemental Fig. 3). And we observed distinct ribosome occupancy patterns across different fractions, which were consistent with previous studies^23,24^ (Supplemental Fig. 4), confirming integrity of our fractionated translational states. In addition, gene abundance measured by SPARP-seq showed a strong correlation with the ribosome-protected fragment counts from these validated Ribo-seq libraries across monosome and polysome fractions (Supplemental Fig. 5). This demonstrates that SPARP-seq reliably quantifies ribosome-associated mRNAs and serves as an accurate proxy for genome-wide translational activity.

### Distinct poly(A) tail length pattern during translation process

Subsequently, we performed detailed analysis of poly(A) tail length distributions across distinct translation fractions using SPARP-seq. The analysis of the overall mRNA population revealed that poly(A) tail length distributions of monosome-associated transcripts were similar to those from polysome fractions, albeit with a distinct peak at ∼60 nt for the monosome fraction (Fig. 1b), confirming previous observations of broad similarity among translation fractions^4^. However, in contrast to this global view, gene-specific analysis revealed that the longest tails were observed on mRNAs from the monosome fraction, and they systematically decreased in length as transcripts associated with heavier polysomes, such as *M6PR* (Fig. 1c-d). Pairwise comparisons at the single-gene level confirmed this trend, showing that a majority of genes exhibited significantly longer median poly(A) tails when associated with monosome compared to their polysome-associated counterparts, with the differences becoming increasingly pronounced as ribosome occupancy increased (Supplemental Fig. 6), implying a dynamic shortening of the poly(A) tail during actively translational status.

Moreover, leveraging the substantial sequencing depth of SPARP-seq, we assessed poly(A) tail length distributions for individual genes with high resolution (Fig. 1e, Supplemental Fig. 7). We observed considerable heterogeneity of poly(A) tail lengths in monosome-associated genes, with many displaying either very short (10–20 nucleotides) or very long tails (exceeding 400 nucleotides). In contrast, the population of polysome-associated mRNAs was far more uniform, with nearly all poly(A) tails falling within a narrower, intermediate window of 50 to 200 nt (Fig. 1e, Supplemental Fig. 7). The observation aligned well with prior studies demonstrating that medium-sized poly(A) tails are optimal for efficient translation^4,5,35,36^.

### Monosome as a Hub for mRNAs with Distinct Poly(A) Tail Signatures and Translational Fates

The observation that poly(A) tails are longest on monosomes, the entry point for translation, and shorten thereafter suggests the monosome acts as a key regulatory hub. We therefore investigated whether different classes of mRNA exhibit distinct poly(A) tail signatures at the monosome stage, reflecting divergent translational fates. We examined the relationship between median poly(A) tail length and gene expression levels. Indeed, we found a striking inverse relationship between median poly(A) tail length and gene expression, where highly expressed genes exhibited shorter poly(A) tails in monosome fraction, explaining the overall sharp peak observed in monosome (Fig. 2a, Supplemental Fig. 8). Conversely, genes in the monosome and polysome fractions exhibiting relatively longer median poly(A) tails were expressed at low levels (Fig. 2a, Supplemental Fig. 8).

**Fig. 2.**
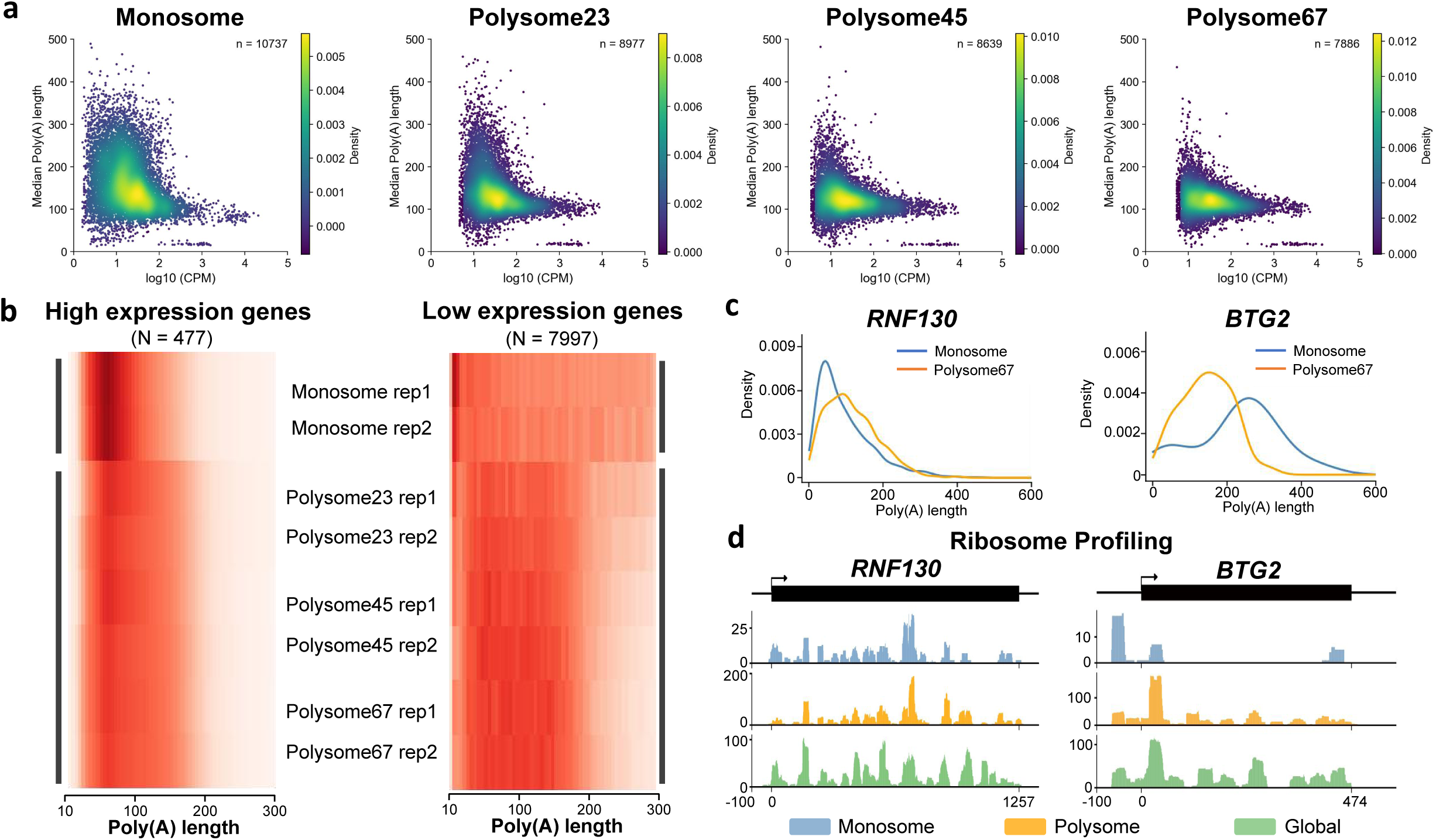
Two models of translational status and poly(A) tail length based on gene expression. **a.** Scatter density plots illustrating the relationship between gene expression level, represented as log10 (CPM), and median poly(A) tail length for biological replicate 1. Plots are shown for each translational fraction (Monosome, Polysome23, Polysome45, Polysome67), with color intensity indicating the density of genes. **b.** Heatmaps comparing the poly(A) tail distribution of highly expressed housekeeping genes (CPM > 200, N = 477) and lowly expressed genes (CPM < 10, N = 7997). Distributions are shown for each ribosomal fraction, with poly(A) lengths grouped (5nt bin). **c.** Kernel density plots comparing the poly(A) tail distribution for highly expressed gene (*RNF130*) and lowly expressed gene (*BTG2*) in monosome versus polysome67 fractions. **d.** Single-gene read coverage plots for highly expressed gene (*RNF130*) and lowly expressed gene (*BTG2*). The data shows read coverage in monosome, polysome, and global ribosome footprinting libraries. Spliced exons were used as the gene reference.

The observation of distinct poly(A) tail patterns between genes of varying abundance prompted us to investigate their translational dynamics at a gene-specific level. We classified genes in monosome into high-expression (Counts Per Million, CPM > 200) and low-expression (CPM < 10) groups and discovered two fundamentally different regulatory strategies. High-abundance genes employ a “workhorse” strategy for constitutive expression. Different from the result of overall gene level trend, these genes displayed short poly(A) tails when associated with monosome, but comparatively longer tails in polysome, e.g., *RNF130* (Fig. 2b-c). This observation offers a new perspective on previous studies that have established short poly(A) tails as a hallmark of highly expressed, efficiently translated genes^5^. It suggests that the correlation may be driven primarily by transcripts in the monosome fraction. To test this, we analyzed Ribo-seq data of monosome, polysome and global, which revealed continuous ribosome coverage across the open reading frames (ORFs) of these monosome-associated mRNAs (Fig. 2d, Supplemental Fig. 9). This confirms that high-abundance genes maintain robust translational capacity even on monosomes, ensuring consistent protein synthesis.

In contrast, low-abundance genes follow a “timer” strategy to maintain a state of translational readiness. These genes exhibited long poly(A) tails on monosome-associated mRNAs that were subsequently shortened upon engagement with polysomes, e.g., *BTG2* (Fig. 2b-c). This pattern, coupled with lower ribosome occupancy in monosome (Fig. 2d, Supplemental Fig. 9), indicates a preparatory state characterized by limited translation. These transcripts appear to be primed by long tails for a later transition into active translation within polysomes, a process accompanied by translation-dependent deadenylation. Together, our results suggested that gene expression level is associated with an mRNA’s translational strategy. Cells appear to utilize two primary strategies: constitutive “workhorse” production for essential genes and a “timer” mechanism for regulated, on-demand protein synthesis.

### Coordinated Regulation of Translation by Alternative Splicing and Poly(A) Tail Length

The translation of distinct splice variants represents an important mechanism for expanding proteome diversity^37–39^, studying translation status of different isoforms via Ribo-seq remains difficult due to the short read length of NGS and often requires large volumes of data with specialized analysis pipeline^40,41^. Taking advantage of the full-length reads in SPARP-seq library, we can easily distinguish mRNA into intron-retained and fully spliced transcripts for individual genes and compare their median poly(A) tail lengths across monosome and polysome fractions (Fig. 3a-b, Supplemental Fig. 10). We found that transcripts retaining introns possessed markedly longer poly(A) tails and were significantly enriched in the monosome fraction compared to their fully spliced counterparts from the same gene (Fig. 3a-b, Supplemental Fig. 10). This result aligns with previous study that ribosome footprints within introns are predominantly linked to monosome^23^. In addition, as the number of ribosome increases, the difference of poly(A) tail length between intron-retained isoforms and fully-spliced isoforms diminished (Fig. 3b, Supplemental Fig. 11). Together, this suggests that the monosome also functions as a quality control checkpoint, where transcripts with retained introns are marked with long poly(A) tails and largely sequestered from entering active, high-throughput translation, likely for clearance by pathways such as nonsense-mediated decay (NMD).

**Fig. 3.**
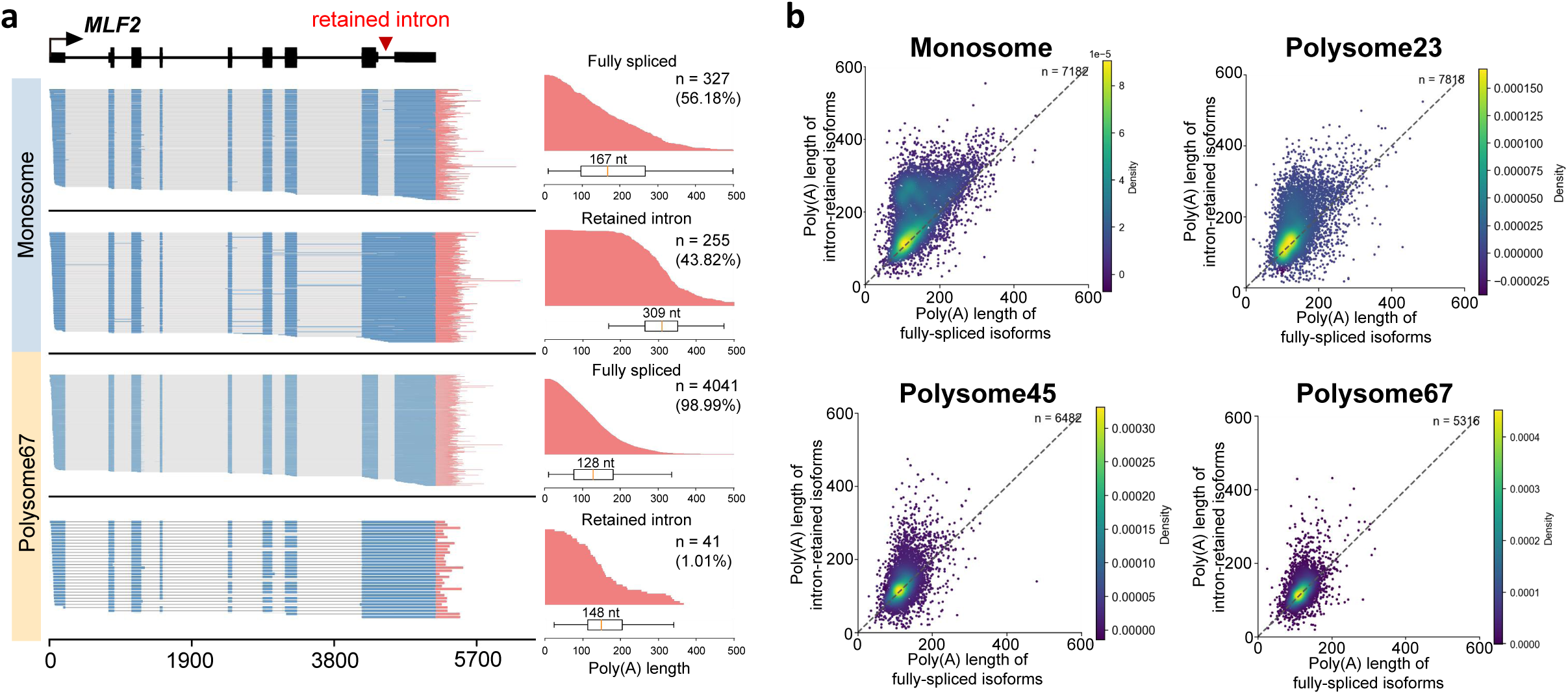
SPARP-seq profiles poly(A) tail length dynamics across transcript Isoforms. **a.** Example of isoform-specific analysis for the *MLF2* gene in the monosome and polysome67 fractions. The left panel displays single-molecule reads aligned to fully spliced or intron-retained isoforms. The right panels show the corresponding poly(A) tail length distributions for each isoform. **b.** Density plots comparing the poly(A) tail lengths of fully-spliced (x-axis) and intron-retained (y-axis) transcript isoforms across all translational fractions. Data from two biological replicates were merged, and only genes with more than 10 reads for both fully-spliced and intron-retained isoforms were included for analysis.

### Increased Poly(A) Tail Length Following Translation Inhibition

To investigate the association between poly(A) tail length and the translation process, we treated 293T cells with the translation inhibitor Torin1^42^ for 0, 2, and 12 hours and performed SPARP-seq analysis. The translation fractions were also separated into monosome, polysome23, polysome45 and polysome67. Treatment for 2 and 12 hours induced a marked shift from polysome to monosome according to the result of sucrose gradient (Fig. 4a), indicating a progressive defect in translation.

**Fig. 4.**
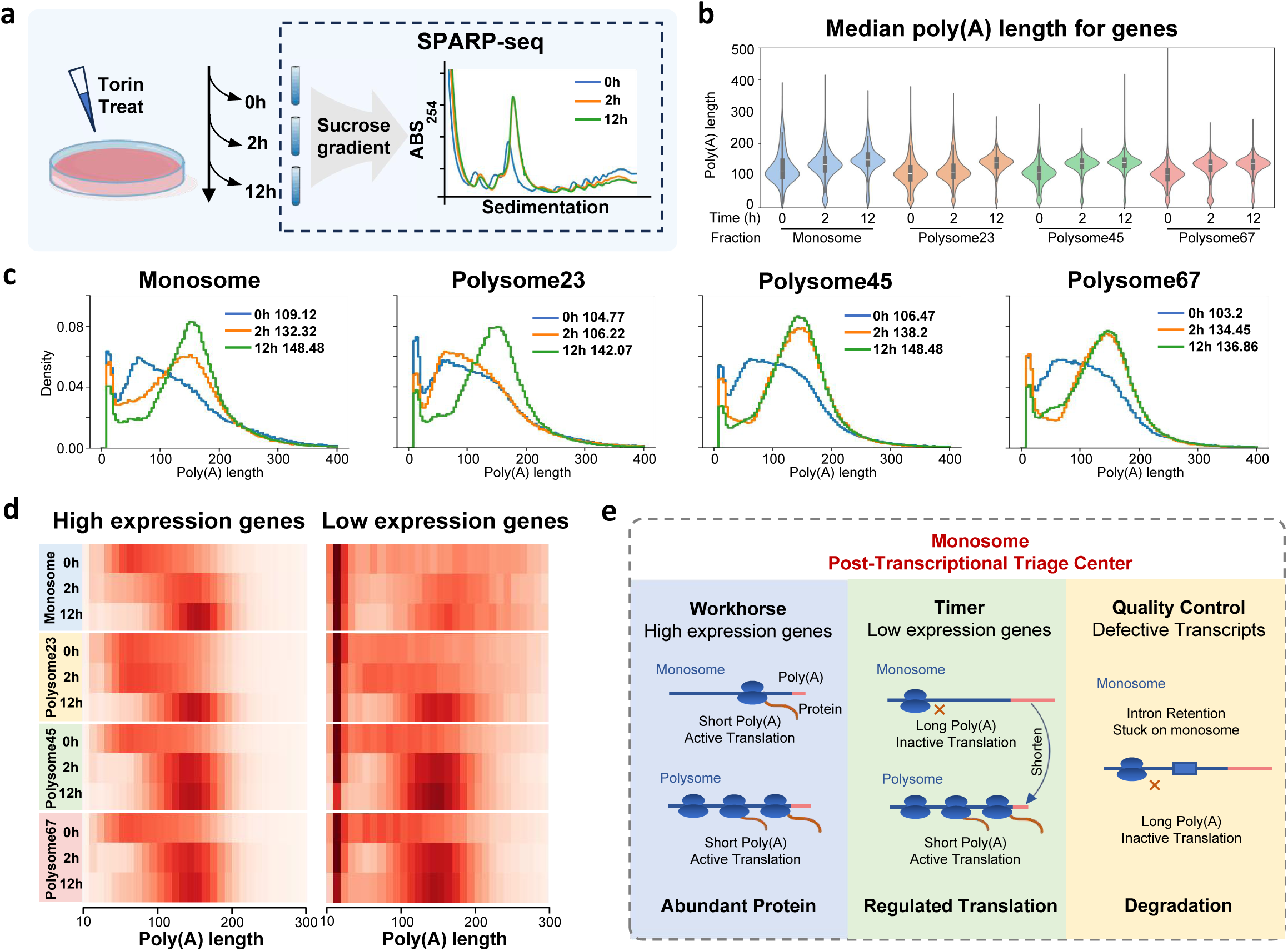
Inhibition of translation by Torin1 leads to global poly(A) tail lengthening. **a.** Schematic of the experimental design of translation inhibition. Cells were treated with Torin1 for 0, 2, and 12 hours. The corresponding sucrose gradient profiles show a time-dependent increase in the monosome peak and a decrease in polysomes after treatment. **b.** Violin plots of the median poly(A) tail length at the gene level in control (0h) and Torin1-treated (2h, 12h) cells for each translational fraction. Genes with more than 10 uniquely mapped poly(A) reads were included. **c.** Overall poly(A) tail length distributions for all reads in each translational fraction at 0, 2, and 12 hours of Torin1 treatment. The plots show a progressive lengthening of poly(A) tails upon translation inhibition. Only reads with poly(A) tails longer than 10 nt were analyzed. **d.** Heatmaps comparing poly(A) tail distribution for highly expressed (CPM > 200) and lowly expressed (CPM < 10) genes after 0, 2, and 12 hours of Torin1 treatment. Four separated translational fractions were shown, with poly(A) length grouped in 10 nt bins. **e.** A summary of monosome-mediated translational regulation. Three primary strategies are shown.

Our analysis from SPARP-seq data revealed that inhibiting translation led to poly(A) tail lengthening across all fractions, but with markedly different kinetics depending on ribosome occupancy. The most immediate and dramatic effect was observed on actively translated mRNAs in heavy polysome fractions (polysome45 and polysome67), where tails lengthened considerably at 2 hours with little subsequent change. In contrast, poly(A) tails of the monosome-associated mRNAs showed a slow, progressive elongation over the 12-hour course. Light polysome (polysome23) exhibited an intermediate phenotype, with only a slight increase at 2 hours followed by a substantial increase at 12 hours (Fig. 4b-c). Moreover, this general lengthening trend was observed in both high- and low-abundance genes (Fig. 4d), further illustrating that changes in poly(A) tail are a key signature of an mRNA’s prior translational status.

Overall, the result illustrated that poly(A) tail length is not a static feature but a dynamic indicator of an mRNA’s real-time translational status, governed by the tight coupling of translation with co-translational deadenylation^43–45^. Blocking this process of translation uncouples mRNA from tail shortening, leading to a net increase in poly(A) tail length. Therefore, poly(A) tail dynamics is not merely a passive consequence of translation, but an active regulatory hub used by the cell to manage the stability and translational competence of its transcriptome in response to environmental cues.

## Discussion

Our study resolves a long-standing paradox in gene expression by providing a high-resolution view of the dynamic interplay between poly(A) tail length and an mRNA’s translational fate. By developing SPARP-seq, we have overcome previous technological limitations that averaged out distinct regulatory programs. The central finding of this work is the establishment of a new model for post-transcriptional regulation, in which the 80S monosome acts as a “post-transcriptional triage center”. Far from being a simple or inactive intermediate, the monosome is a dynamic hub where the translational trajectory of an mRNA is decoded from its poly(A) tail signature.

Our data revealed at least two distinct strategies for translational control that are integrated at the level of the poly(A) tail and sorted at the monosome. For highly expressed housekeeping genes, an intrinsically short and stable poly(A) tail appears to license efficient, constitutive translation directly from monosomes. This “workhorse” strategy, which supports previous findings that monosomes can be translationally active, may confer a distinct cellular advantage. This mode of synthesis could provide a steady, reliable supply of essential proteins while avoiding the potential for ribosomal traffic jams and the sequestration of translational resources associated with the formation of large polysomes. In contrast, for many regulated and lower-abundance genes, a long poly(A) tail on monosome-associated transcripts appear to act as a “timer”, priming the transcript for a robust burst of protein synthesis that is coupled to progressive, translation-dependent deadenylation upon recruitment to polysomes. In summary, our SPARP-seq data demonstrated that both short and long poly(A) tails can be associated with translation, but for different classes of genes with distinct functional requirements.

Furthermore, the monosome also functions as a critical quality control checkpoint^23,46^. We found that translationally repressed transcripts, such as those retaining introns^23,47^, are preferentially enriched on monosomes and marked with long poly(A) tails. This enrichment likely either routes them for degradation through pathways like nonsense-mediated decay (NMD) or keeps them in a dormant state, awaiting a specific signaling or environmental cue to activate translation^48,49^. Together, these findings paint a model of the monosome as a sophisticated sorting station that integrates an mRNA’s identity (e.g., housekeeping, regulated, or defective) with its poly(A) tail status to orchestrate its fate (Fig. 4e).

The tight coupling of active translation with deadenylation was functionally confirmed by our Torin1 inhibition experiments, which led to a global lengthening of poly(A) tails as ribosomes were stalled. While our data strongly suggests this is primarily due to the uncoupling of translation from co-translational deadenylation, we cannot exclude an additional contribution from mTOR-dependent regulation of the polyadenylation and deadenylation machinery itself, a possibility that warrants future investigation.

## Experimental Procedures

### Cell culture and mTOR inhibition treatment

HEK293T cells were maintained in DMEM (Gibco, 11965092) supplemented with 10% FBS (Cellmax, SA301.02) and 1% Penicillin-Streptomycin (Gibco, 15070063) in a humidified atmosphere containing 5% CO_2_ at 37°C. Cells were subcultured the day before harvesting to ensure confluency reached approximately 70% at the time of collection. For translation inhibition experiments, cells were treated with either a vehicle control or 250 nM Torin1 for 0, 2, and 12 hours prior to harvesting.

### Polysome profiling

Polysome profiling was performed as previously described^4,50^. Briefly, cultured cells were treated with cycloheximide at a final concentration of 100 µg/mL and incubated for 10 minutes at 37°C to arrest ribosomes on translated mRNA. After treatment, cells were immediately placed on ice and washed twice with ice-cold PBS containing 100 µg/mL cycloheximide to remove residual media. Cells were then gently scraped from the plate using a cell scraper and transferred to a 15 mL centrifuge tube. The harvested cells were centrifuged at 1,000 × *g* for 5 minutes at 4 °C, and the supernatant was discarded. The resulting cell pellet was resuspended in 300 µL lysis buffer containing 10 mM HEPEs pH 7.4, 5 mM MgCl2, 150 mM KCl, 1% NP40, 1mM DTT, 100 ug/mL cycloheximide, 1x protease inhibitor cocktail (EDTA-free), 200 units/mL of RNase inhibitor (40 U/μL, Vazyme, R301-03). The suspension was lysed by passing through a 25-gauge needle 10 times, followed by incubation on ice for 30 minutes to ensure complete lysis. The lysate was centrifuged at 1,000 × *g* for 5 minutes at 4°C to drop cell nuclei and large debris. The supernatant was carefully transferred to a fresh tube and centrifuged again at 16,363 × *g* for 10 minutes at 4 °C to remove smaller debris. The resulting clear supernatant was transferred to a new tube and kept on ice. To isolate total RNA, 20 µL lysate was transferred to a new tube and mixed with 300 µL of TRIzol LS reagent.

10% and 50% sucrose buffer were prepared in a buffer containing 10 mM HEPES pH 7.4, 5 mM MgCl₂, and 150 mM KCl, followed by filtration through a 0.22 µm filter to ensure sterility. A 10%-50% sucrose gradient was prepared using a Biocomp Gradient Master. The cell lysis supernatant was carefully loaded onto the prepared gradient sucrose and centrifuged at 36,000 rpm using Beckman SW-41 Ti rotor for 150 min at 4 °C. Following centrifugation, the gradients were fractionated using Biocomp Gradient Fractionator with absorbance monitored at 254 nm to detect ribosomal subunits and polysomes. Fractions corresponding to monosome and polysomes, including groups containing 2–3, 4–5, and 6–7 ribosomes, were pooled separately. These pooled fractions were used for downstream RNA extraction.

### RNA extraction

RNA extraction was performed using TRIzol LS reagent. Briefly, 3 volumes of TRIzol LS were added to each sample, followed by vortexing for 10 seconds to ensure thorough mixing. The mixture was incubated at room temperature for 5 minutes and then centrifuged at 14,000 × *g* for 10 minutes at 4 °C. The upper aqueous phase was carefully transferred to a new 1.5 mL tube, followed by adding 0.2 × volume of chloroform. The mixture was vortexed for 10 seconds and centrifuged again at 14,000 × *g* for 10 minutes at 4 °C. The upper aqueous phase was collected and subjected to further RNA purification using the RNA Clean & Concentrator Kit (Zymo Research, R1015), following the manufacturer’s instructions. The purified RNA was eluted in 16 μL of nuclease-free water and quantified using the Qubit RNA High Sensitivity Assay.

### SPARP-seq Library construction

#### rRNA depletion and linker ligation

100 ng to 1500 ng of extracted RNA was first subjected to ribosomal RNA depletion using riboPOOL probes targeting human, mouse, and rat rRNA (dp-R096-000050) according to the manufacturer’s protocol. The depleted RNA was cleaned and concentrated to 6 μL using RNA Clean & Concentrator Kit. To capture polyadenylated RNA and non-polyadenylated transcripts, 50 pmol of 3’ end-blocked oligodeoxynucleotide (NEB S1315S) (5’-rAppCTGTAGGCACCATCAAT-NH2 −3’) was ligated to the 3’ end of the rRNA-depleted RNA. The reaction mixture was incubated at 65°C for 5 minutes to facilitate annealing and then cooled on ice for at least 1 minute. Next, the following components were added to the reaction: 2 μL of 10× T4 RNA Ligase Reaction Buffer (NEB, M0242), 10 μL of 50% PEG 8000 (NEB, M0242), 1 μL of murine RNase inhibitor, and 1 μL of T4 RNA Ligase 2, truncated K227Q (NEB, M0242). The reaction mixture was thoroughly mixed by pipetting and incubated at 16°C for 10 hours to ensure efficient ligation.

#### Reverse transcription and PCR amplification

The ligated RNA was purified using ZYMO RNA Clean & Concentrator-5 kit (ZYMO, R1013) and eluted with 9 μL RNase-free H_2_O. To initiate reverse transcription, 1 µL cDNA RT primer (5’-phos/ACTTGCCTGTCGCTCTATCTTCATTGATGGTGCCTACAG -3’, 12 μM) were added to the eluted RNA and incubated at 65 °C for 5 min. The annealed RNA was placed on ice for more than 1 minute and the other components were added containing 4 μL 5x RT Buffer, 1 μL murine RNase inhibitor, 2 μL Strand-Switching Primer (SSP, at 10 µM) and 1 μL Nuclease-free H_2_O. After incubating at 42 °C for 2 minutes, 1 µL of Maxima H Minus Reverse Transcriptase (Thermo Fisher, EP0752) was introduced, bringing the total reaction volume to 20 μL. Reverse transcription was carried out with the following thermal cycling conditions: 42 °C for 90 minutes, followed by 10 cycles of 50 °C for 2 minutes and 42 °C for 2 minutes, and a final step at 85 °C for 5 minutes.

To minimize the PCR bias, the optimal PCR cycle number was determined by assessing the smear pattern on a gel electrophoresis, as previously described^51^. Based on the results, the ideal cycle number was selected for large-scale PCR. Three large-scale PCR reactions were set up using 2 μL of the cDNA product and 0.4 μL of a primer containing a specific barcode (Nanopore SQK-PCB109) per reaction.

### Nanopore sequencing of SPARP-seq Library

Following PCR amplification, the product was treated with 1 μL Exonuclease Ⅰ (NEB, M0293) at 37℃ for 15 min to remove residual single-stranded DNA, followed by heat inactivation at 80℃ for 15 minutes. The PCR product was then purified using 0.8× Ampure XP beads (Beckman, A63880) according to the manufacturer instructions. The purified library was eluted with 15 μL EB buffer. The concentration of the library was quantified using DNA Qubit High Sensitivity Assay. Samples with different barcodes were pooled together for sequencing, with each sample contributing an equal molar amount to achieve a final pooled concentration of 100 fmol. The mixed library was loaded onto FLO-PRO106 or FLO-MIN106 (R9.4.1) flow cells and sequenced on the MinION Mk1B or PromethION 2 Solo for 100 hours. Data collection and sequencing were managed by the MinKNOW software following default parameters.

### SPARP-seq data processing

Raw fast5 files generated from nanopore sequencing were basecalled with Guppy (v4.0.11) with the ‘dna_r9.4.1_450bps_hac.cfg’ model, retaining reads with a mean quality score above 7. Barcode demultiplexing was performed with the parameter ‘--barcode_kits “SQK-PCB109“’, producing separate Fastq files for each barcode. The resulting data were aligned to the hg38 reference genome using minimap2 with the parameters ‘-ax splice --secondary=no’. Adapter identification and poly(A) tail length estimation were performed following the FLEP-seq pipeline^51,52^. Reads uniquely mapped to protein-coding genes were retained for calculating counts per million (CPM). Reads with poly(A) tail lengths longer than 10 nucleotides were selected for poly(A) tail length analysis. For each gene, only those with more than 10 uniquely mapped poly(A) reads were included in calculating the median poly(A) tail length.

To detect retained introns at single-read level, data from the same fraction across two replicates were combined. Reads mapping to the intron region with over 80% coverage were considered retained intron reads, while reads not mapping to the intron region were classified as fully spliced. Gene-level analysis of poly(A) tails on retained intron versus fully-spliced transcripts required a minimum read count of 10 for each category.

### Global, monosome and polysome ribosome footprinting

To characterize the global translational activity, ribosome profiling was performed as previously described^4^. Specifically, the total RNA concentration of cell lysate was measured using the Equalbit RNA BR Assay Kit (Vazyme, EQ212-01). RNase I (Ambion, AM2294) was used for RNA digestion at the ratio of 0.15 U RNase I per microgram total RNA. The reaction was incubated at 25 °C with gentle shaking at 400 rpm for 30 minutes, then 10 µL of SUPERaseIN RNase inhibitor (Invitrogen, AM2694) was added to stop reaction. The monosome-associated particles were separated using a 10%-50% sucrose gradient as above described. RNA extraction was performed following the protocols outlined above.

To characterize translational activity in monosome and polysome, ribosome profiling was performed as previously described^23,24^. Briefly, monosome and polysome fractions were separated using a 10–50% sucrose gradient centrifugation as described above. The separated fractions were processed using Amicon-Ultra 100K columns (Millipore, UFC910024 and UFC810024) to concentrate the samples. For monosome fractions, the columns were centrifuged at 5,000 × g at 4 °C for 20 minutes. For polysome fractions, centrifugation was extended to 30 minutes under the same conditions. The concentrated samples were diluted with 5 ml or 10 ml of lysis buffer containing 100 µg/mL cycloheximide to reduce sucrose concentration. The diluted samples were subjected to another round of centrifugation using the same Amicon-Ultra 100K columns under identical conditions. Finally, the RNA concentration of the processed samples was quantified using the Equalbit RNA BR Assay Kit (Vazyme, EQ212-01).

For RNase digestion, RNase I was added to 15 µg of RNA at the ratio of 20 U RNase I per microgram RNA in 1000 µL lysis buffer. The mixture was incubated at 25 °C with gentle shaking at 400 rpm for 30 min, then 10 µL of SUPERaseIN RNase inhibitor was added to stop the digestion. To aid RNA precipitation, 2 μL of GlycoBlue™ Coprecipitant (15 mg/mL, Thermo Fisher) was added to the reaction. Then, the 500 µL digested product was loaded onto a 600 μL 34% sucrose cushion in a ultracentrifuge tube. Centrifugation was performed at 100,000 rpm for 1 hour at 4 °C using a Beckman TLA-120.2 rotor. Following centrifugation, the supernatant was carefully removed, and the depositing pellet corresponding to monosome was resuspended in 500 μL of TRIzol LS reagent. RNA extraction from the resuspended pellet was performed as described above.

### Ribosome footprinting library construction

The extracted RNA was loaded onto a 15% Urea-PAGE gel, and RNA fragments between 26 and 34 nucleotides were excised. The excised RNA was recovered using the ZR small-RNA PAGE Recovery Kit (R1079). To remove residual ribosomal RNA, the Illumina Ribo-Zero Gold rRNA Removal Kit (Human/Mouse/Rat, 20040525) was used. To prepare RNA for library construction, T4 Polynucleotide Kinase (PNK, NEB, M0201S) was used for 5’ phosphorylation and 3’ dephosphorylation of the RNA fragments. The RNA was subsequently purified again using the Zymo RNA Clean & Concentrator Kit.

The RNA library was constructed following the instruction manual of the NEBNext® Multiplex Small RNA Library Prep Set for Illumina (NEB, E7330S) protocol. The products were purified with DNA clean and concentration kit (Zymo, D4004). And size selection of 160 bp DNA was carried out by electrophoresis through a 6% Polyacrylamide Gel. The final library was sequenced on the Illumina NovaSeq 6000 platform.

### Ribosome footprinting data processing

The analysis pipeline of Ribosome profiling is followed by previous studies^25,53^. Raw reads were generated as paired-end data. Due to the short length of the fragments (∼30 bp) after enzyme digestion, only a single file from the paired-end data was used for analysis. Adapter trimming was performed using fastp to remove 3’ adapter. Bowtie with the parameter ‘-v 2 -p 8’ was used to remove the reads aligned to rRNA. The remaining reads were mapped to the hg38 genome using STAR with the following parameters: --outFilterType BySJout, --runThreadN 16, --outFilterMismatchNmax 2, --outFilterMultimapNmax 1, --outFilterMatchNmin 16, and --alignEndsType EndToEnd. The longest transcript of each gene was used as the reference annotation. Quality control of the ribosome profiling data was performed using RiboseQC to assess three-nucleotide periodicity of ribosome footprint. Based on these analyses, reads with lengths of 25–37 nt were selected as ribosome footprints and used for downstream ribosome profiling analysis.

Filtered reads were further processed to remove residual rRNA and mitochondrial RNA sequences using Bowtie with parameters ‘-v 2 -p 8’. The remaining reads were aligned to the genome using STAR, with the longest transcript of each gene used as the reference annotation. Alignment parameters included --outFilterType BySJout, --runThreadN 16, --outFilterMismatchNmax 2, and --outFilterMultimapNmax 1. Only uniquely mapped reads were retained for downstream analysis. Gene-level read counts were quantified using featureCounts, with reads mapped to exons specified by the parameter -t exon. Reads per kilobase per million mapped reads (RPKM) were then calculated for each gene.

## Accession Numbers

The SPARP-seq data generated in this study have been deposited in the Genome Sequence Archive in National Genomics Data Center, China National Center for Bioinformation / Beijing Institute of Genomics, Chinese Academy of Sciences (GSA: PRJCA035252) that are publicly accessible at https://ngdc.cncb.ac.cn/bioproject/browse/PRJCA035252. For reviewers, a temporary access to the data is available through the following link: https://ngdc.cncb.ac.cn/gsa-human/s/9Mk1U0F3.

## Code availability

Source code for analysis is available at https://github.com/ZhaiLab-SUSTech/SPARP-seq.

## Acknowledgments

The group of J.Z. is supported by the Biological Breeding-National Science and Technology Major Project (2023ZD04073), National Natural Science Foundation of China (32325031, 32470601), the Program for Guangdong Introducing Innovative and Entrepreneurial Teams (2016ZT06S172), National Natural Science Foundation of China (32222039).

## Author Contributions

J.Z. and Y.S. conceived and designed the experiments. Y.S., Z.S., Y.L. performed the experiments. Y.S. and J.L. analyzed the data. S.L., X.C. and W.C provided materials and conceptual insights. J.Z. oversaw the study. Y.S. and J.Z. wrote the manuscript, and all authors revised the manuscript.

---

**Supplementary Fig. 1.**
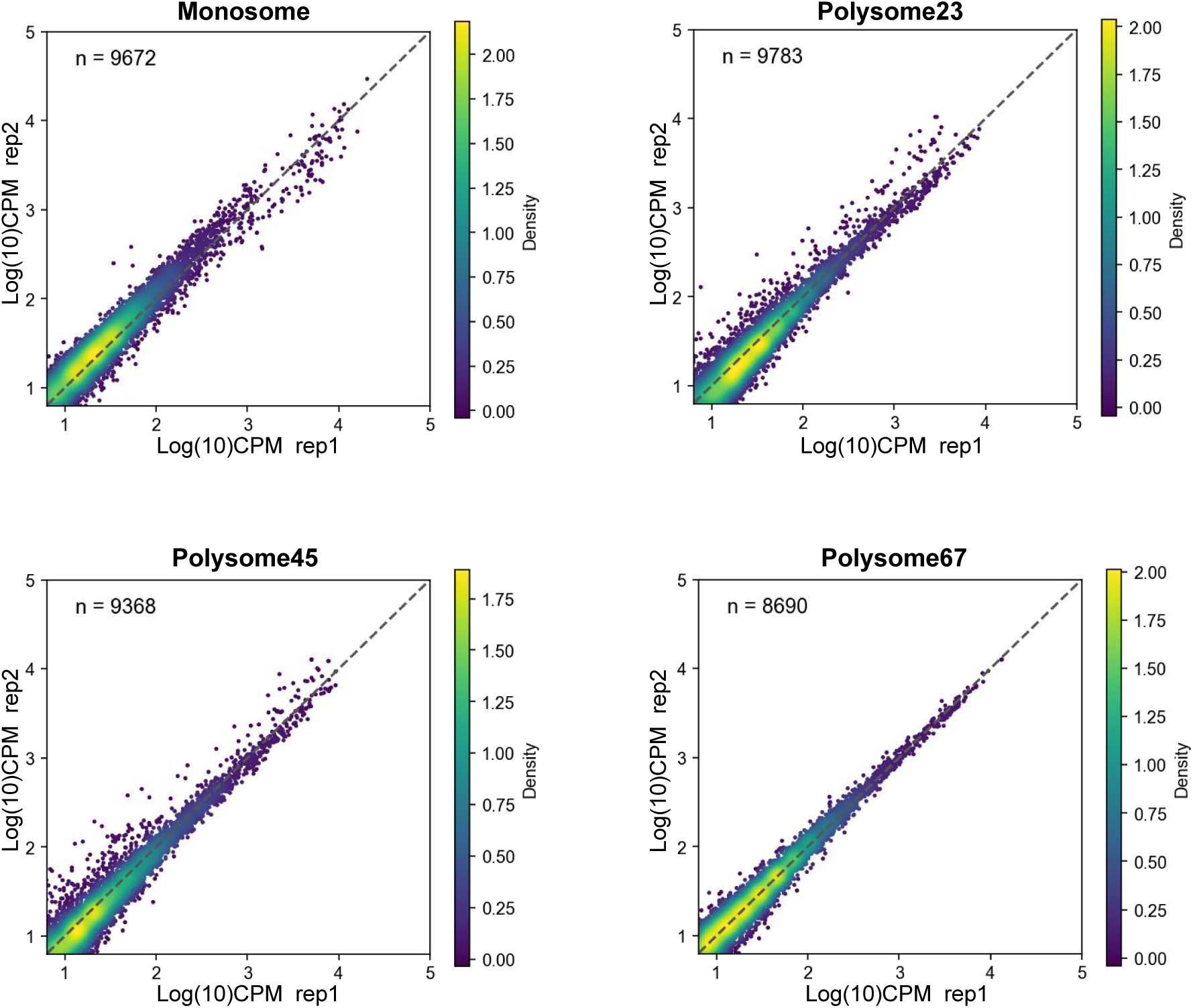
Scatter plots showing the association of gene read counts between biological replicates (Replicate 1 vs. Replicate 2) across monosome, polysome23, polysome45, and polysome67 fractions. Only genes with mapped read counts more than 10 in both replicates were included.

**Supplementary Fig. 2.**
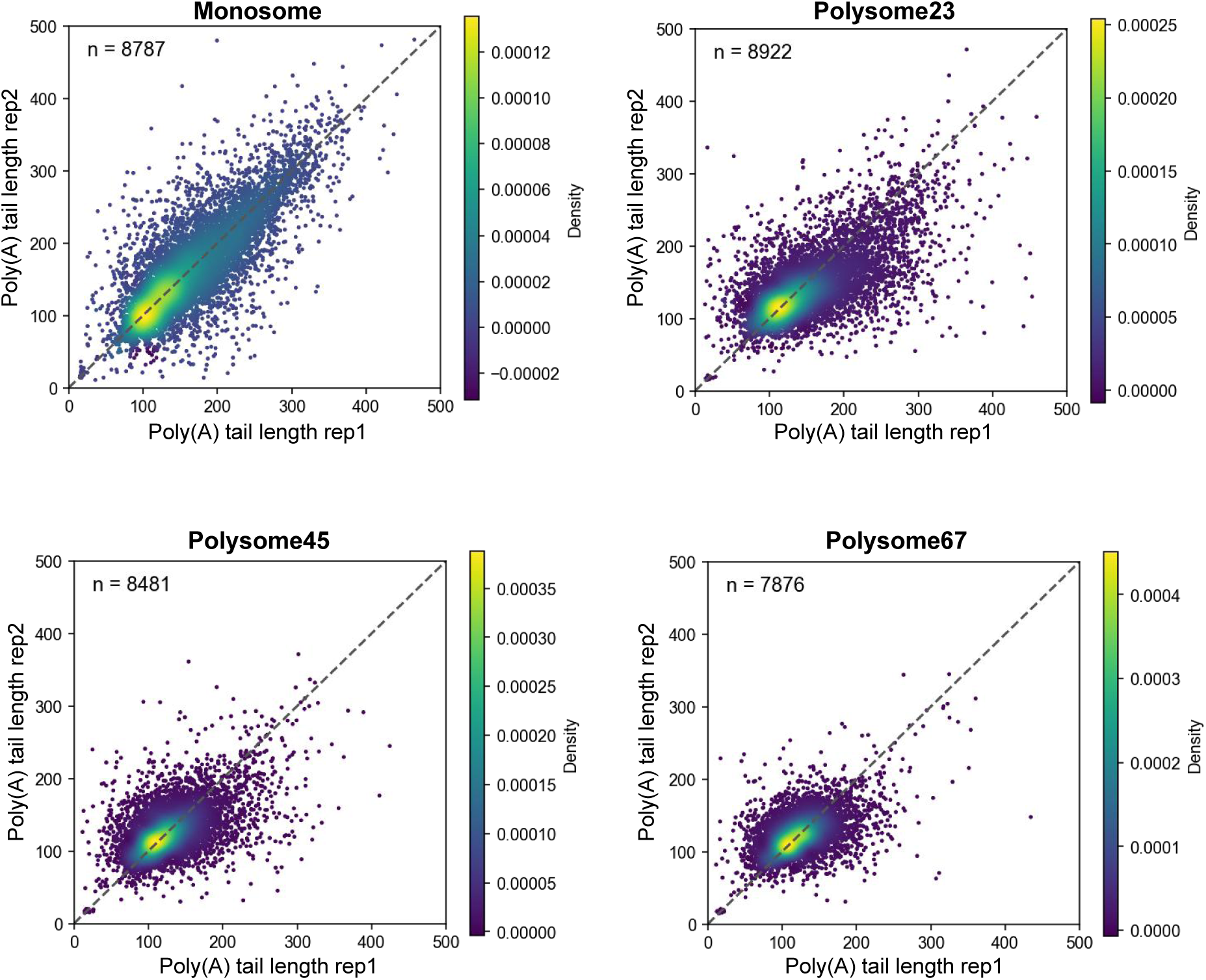
Scatter plots showing the association of gene median poly(A) tail lengths between biological replicates (Replicate 1 vs. Replicate 2) across monosome, polysome23, polysome45, and polysome67 fractions. Only poly(A) tail lengths longer than 10 nt were considered as poly(A) reads, genes with poly(A) reads count more than 10 in both replicates were included.

**Supplementary Fig. 3.**
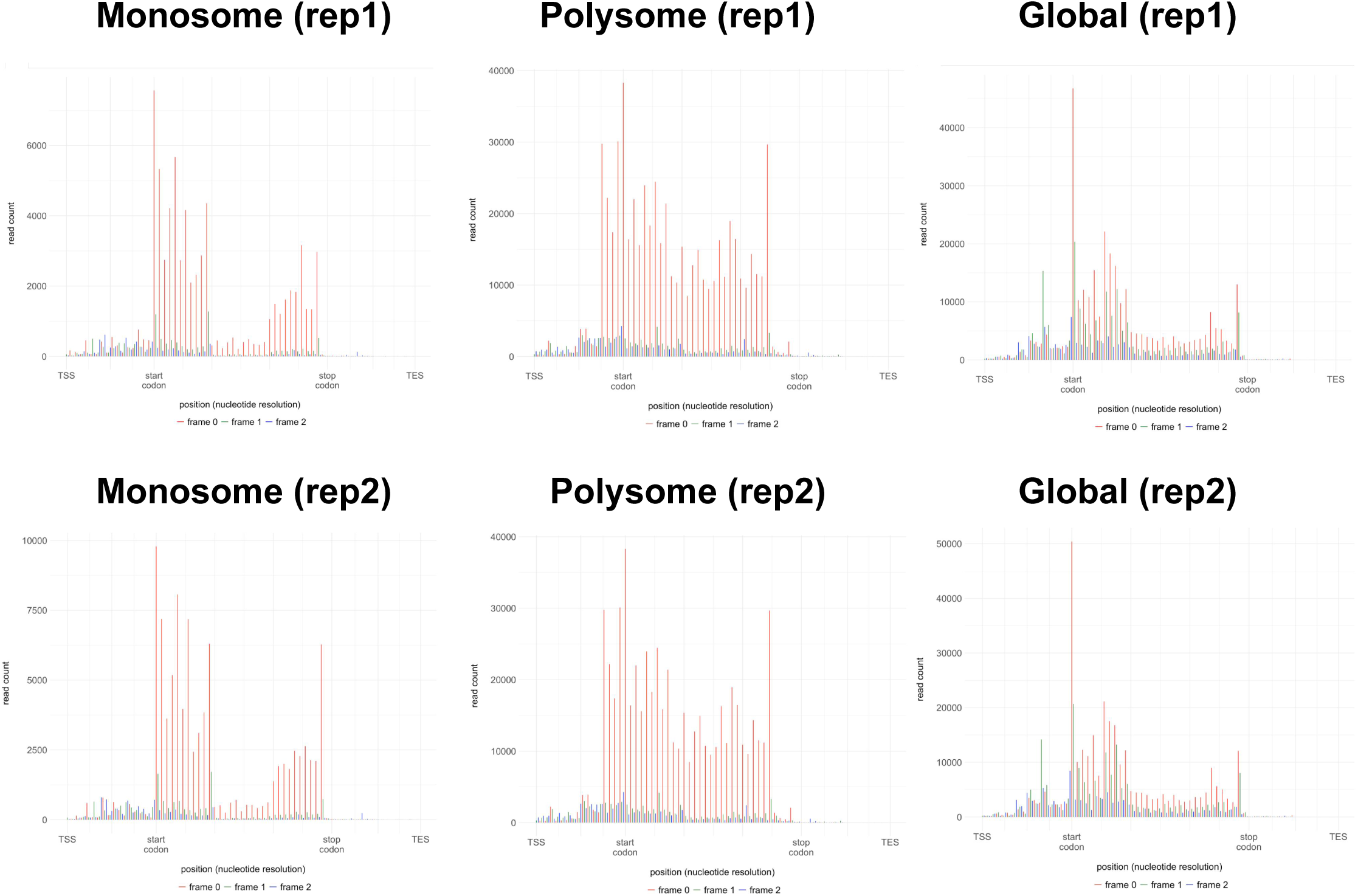
Aggregation plots of ribosome profiling data showing the distribution of the 5’ end positions of reads within the gene region.

**Supplementary Fig. 4.**
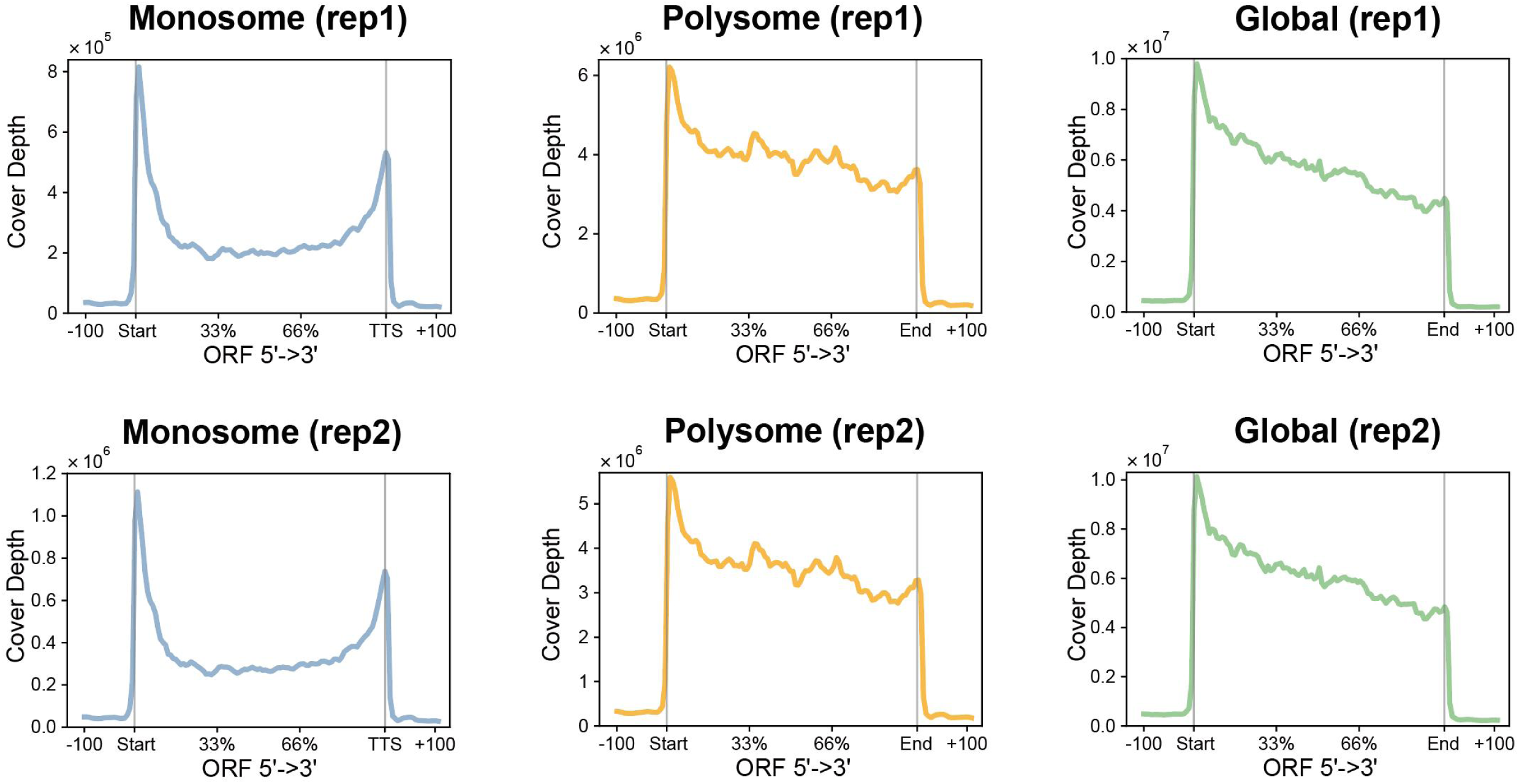
Metagene plots of 25-37 nt fragments throughout the transcript ORF in monosome, polysome and global Ribo-seq, including 100 bp upstream and downstream extensions of the ORF region.

**Supplementary Fig. 5.**
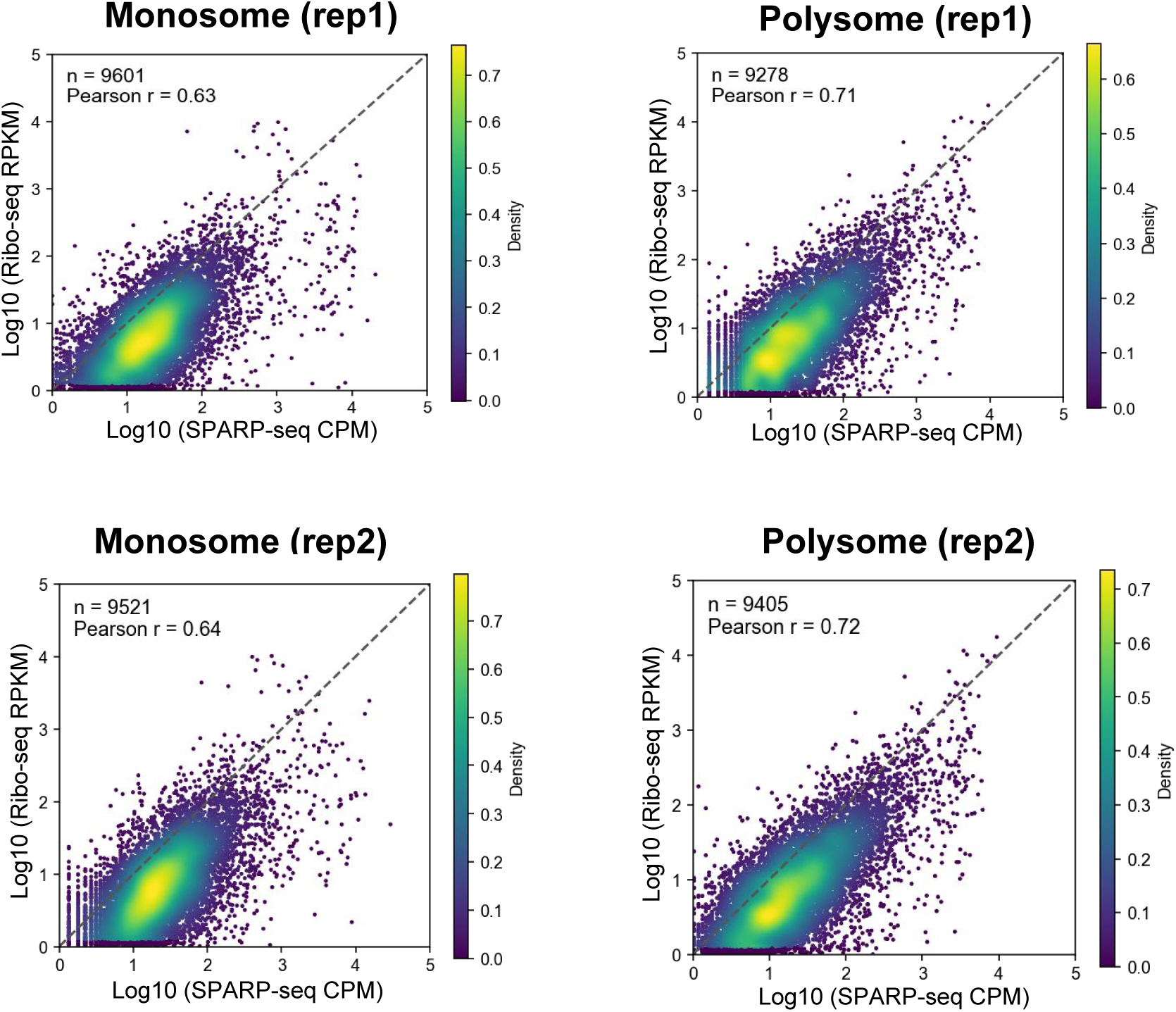
Scatter density plots showing the association of gene abundance between SPARP-seq (Log10 CPM) and Ribo-seq (Log10 RPKM) for rep2, with color intensity representing data density. The polysome Ribo-seq result was compared with the polysome67 result of SPARP-seq libraries. Only Genes with more than 10 mapped reads in SPARP-seq and Ribo-seq were considered.

**Supplementary Fig. 6.**
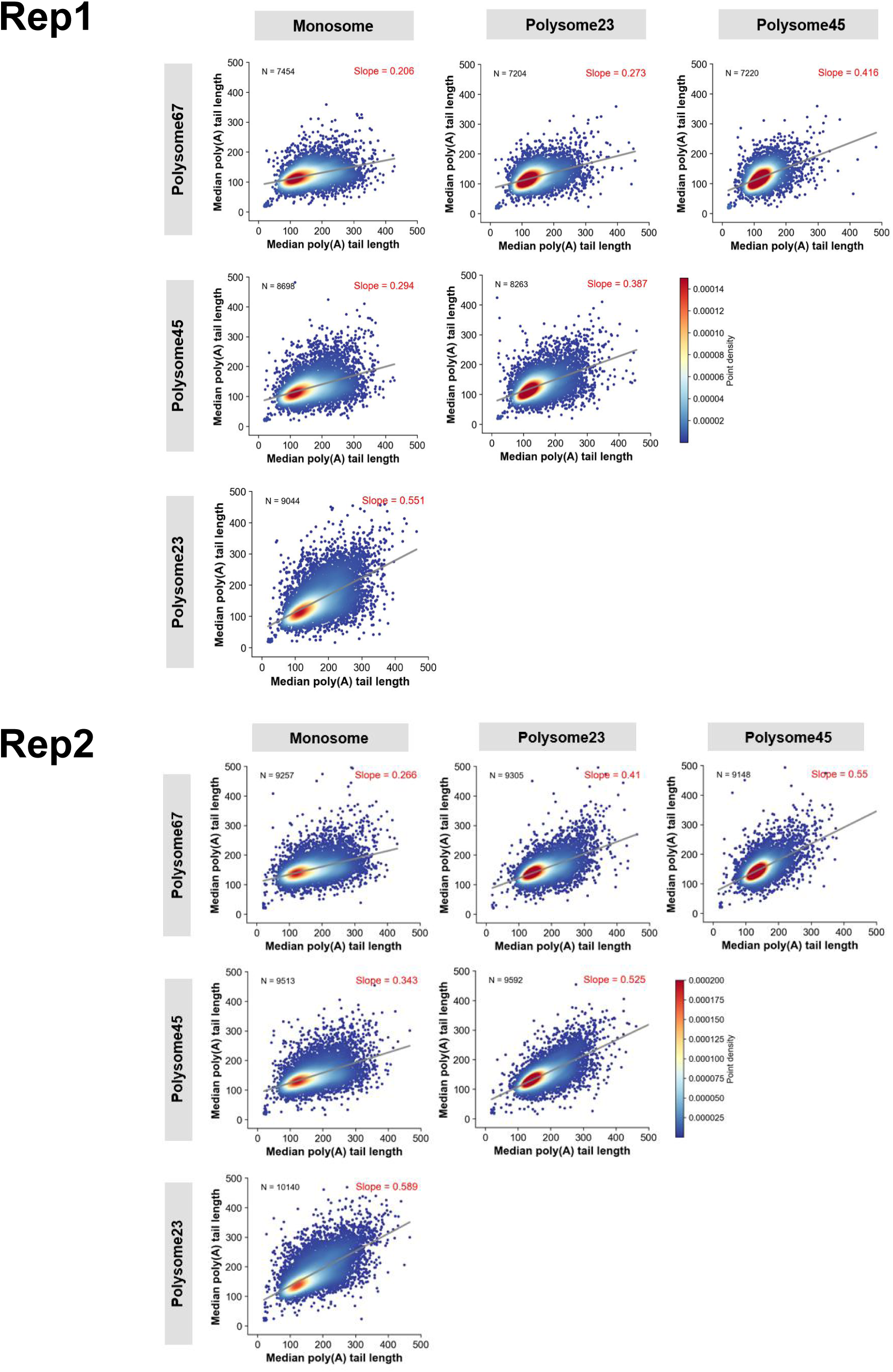
Pairwise comparison of gene median poly(A) tail length among libraries for replicate1 and replicate2. Only genes with more than 10 mapped reads were used for analysis. Reads with poly(A) tail length longer than 10 were considered for calculating the median poly(A) tail length.

**Supplementary Fig. 7.**
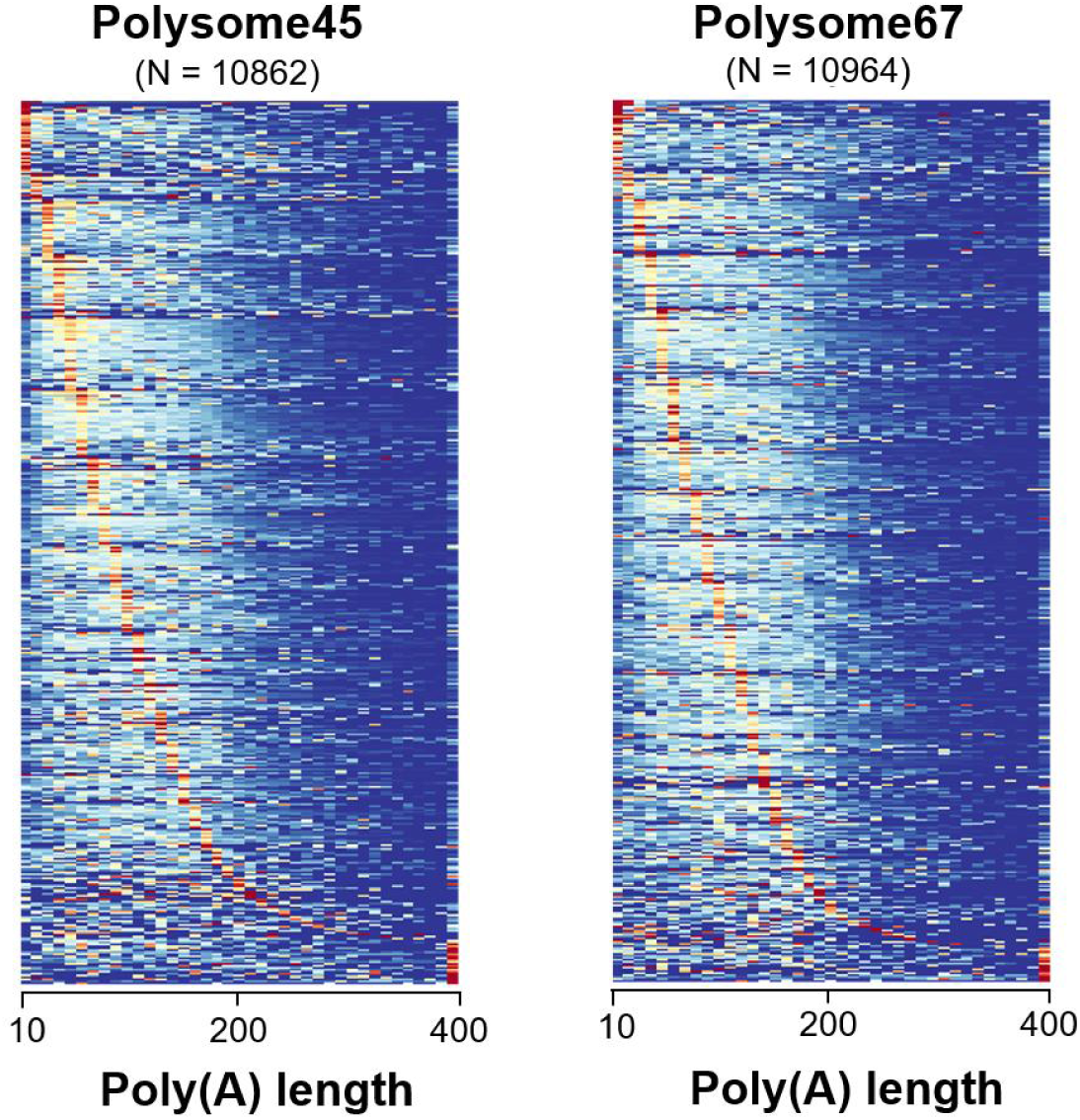
Heatmap showing poly(A) tail length pattern of all genes in SPARP-seq (polysome23 and polysome45) libraries. Data from two biological replicates within each fraction were merged. Each row in the plot represents the poly(A) tail length distribution of a single gene (10 nt bin).

**Supplementary Fig. 8.**
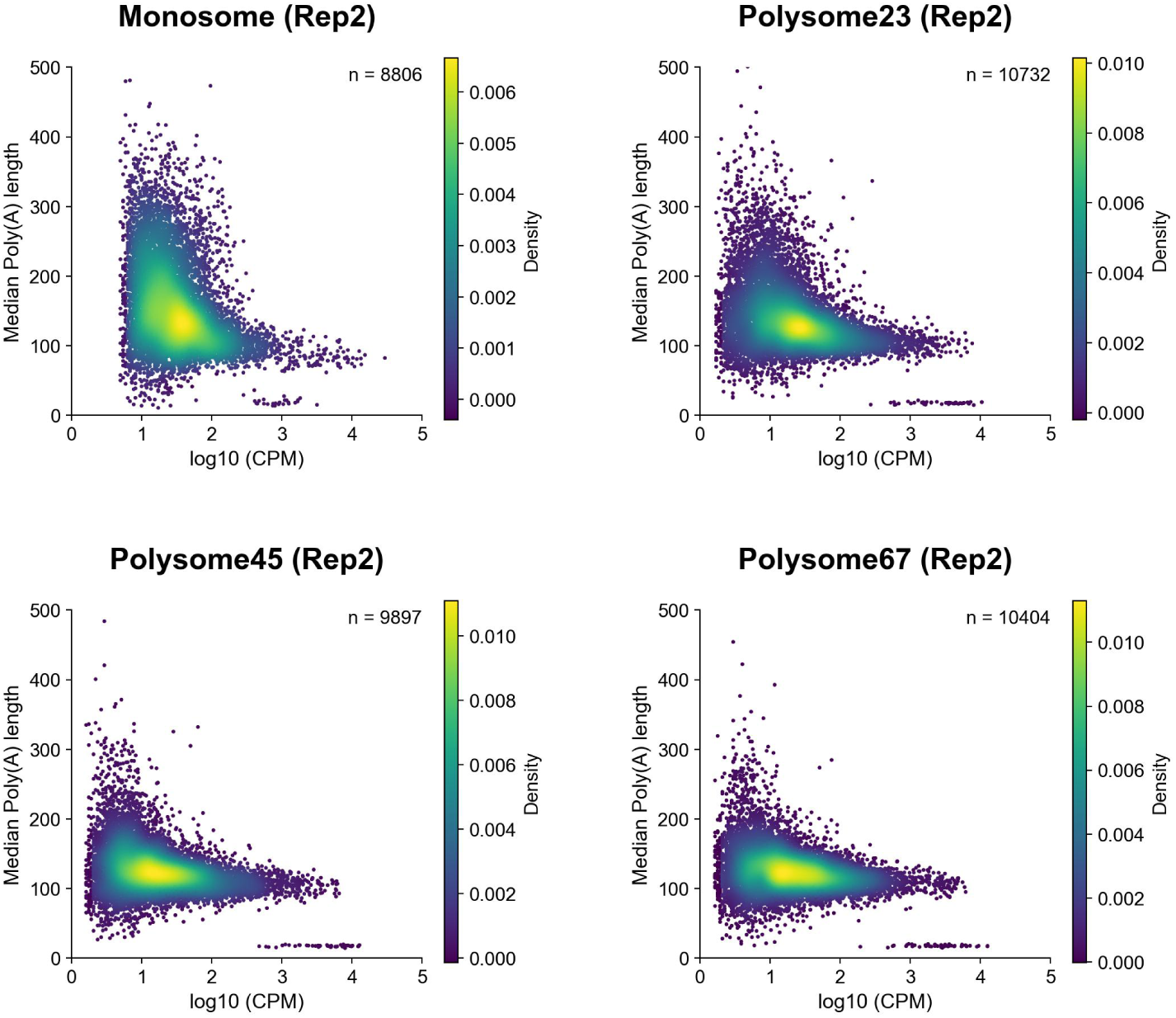
Scatter density plots showing the relationship between gene abundance (Log10 CPM) and median poly(A) tail lengths in the SPARP-seq biological replicate 2 dataset, with color intensity representing gene density. Only Genes with more than 10 mapped reads and reads with poly(A) tail length longer than 10 nt were considered.

**Supplementary Fig. 9.**
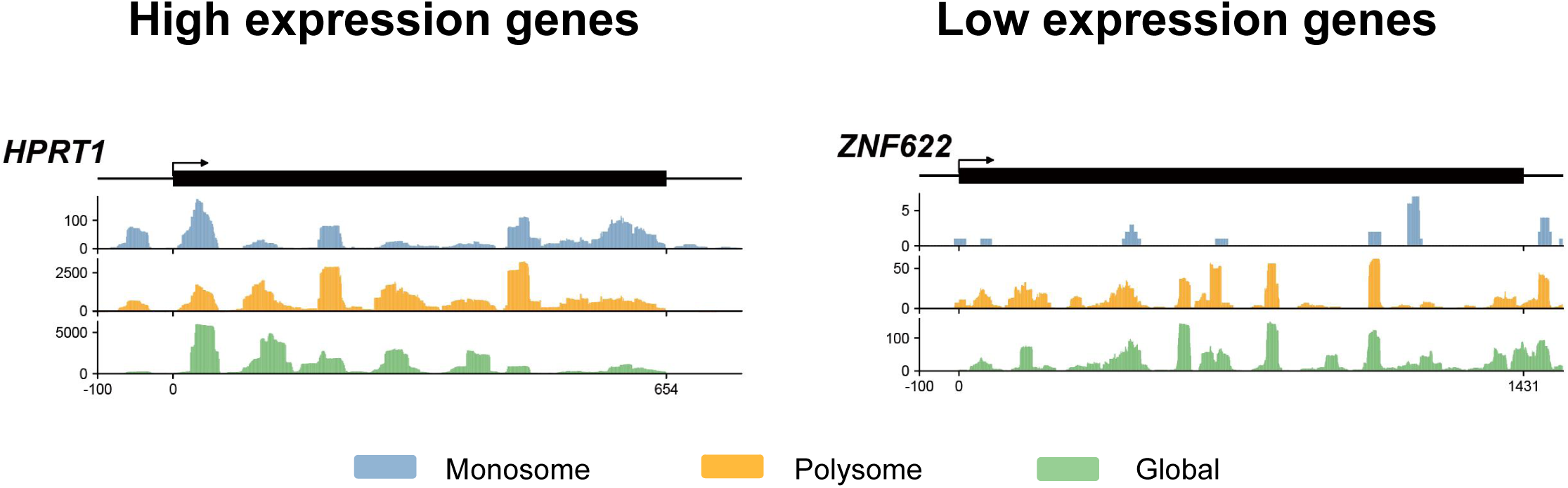
Single-gene read coverage of high expression gene *HPRT1* (left) and low expression genes *ZNF622* (right) in monosome, polysome and global ribosome footprinting libraries. Exons were extracted and spliced to serve as the gene reference.

**Supplementary Fig. 10.**
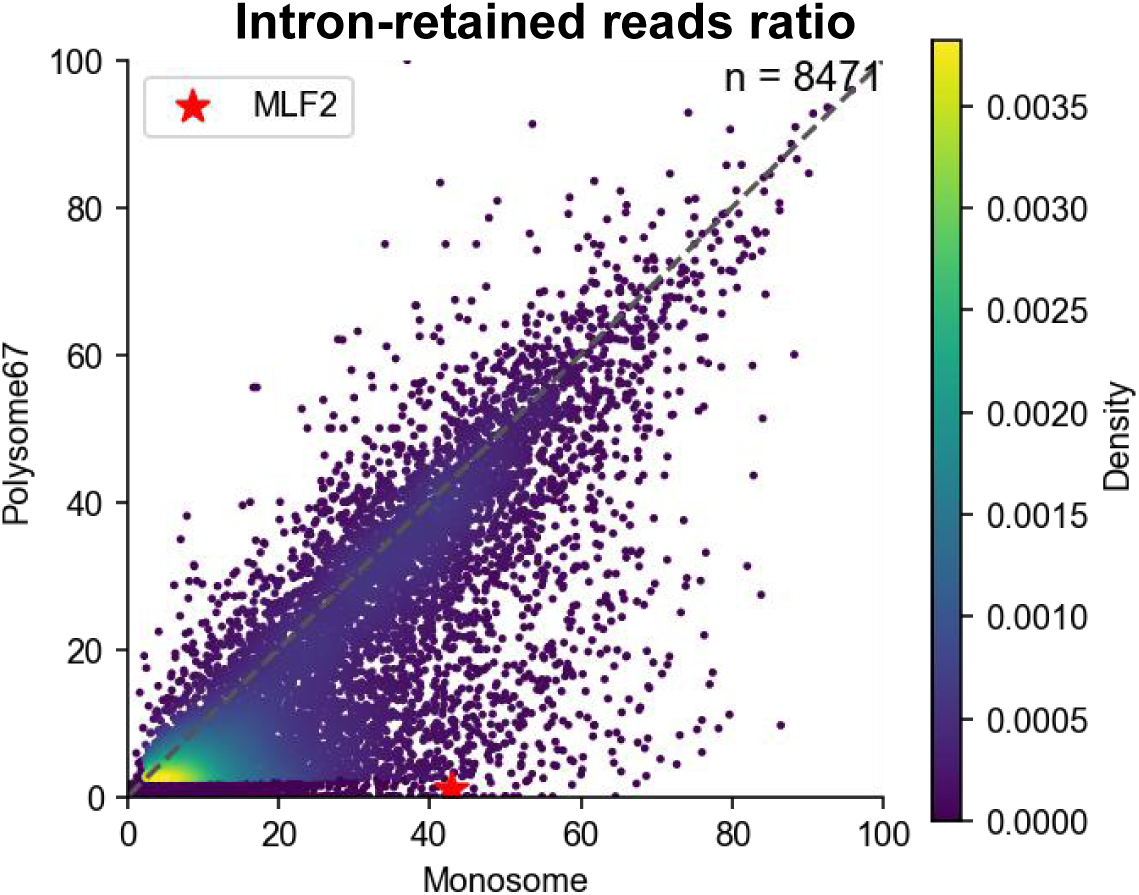
Comparison of intron-retained reads ratio to total reads for each gene between monosome and polysome67.

**Supplementary Fig. 11.**
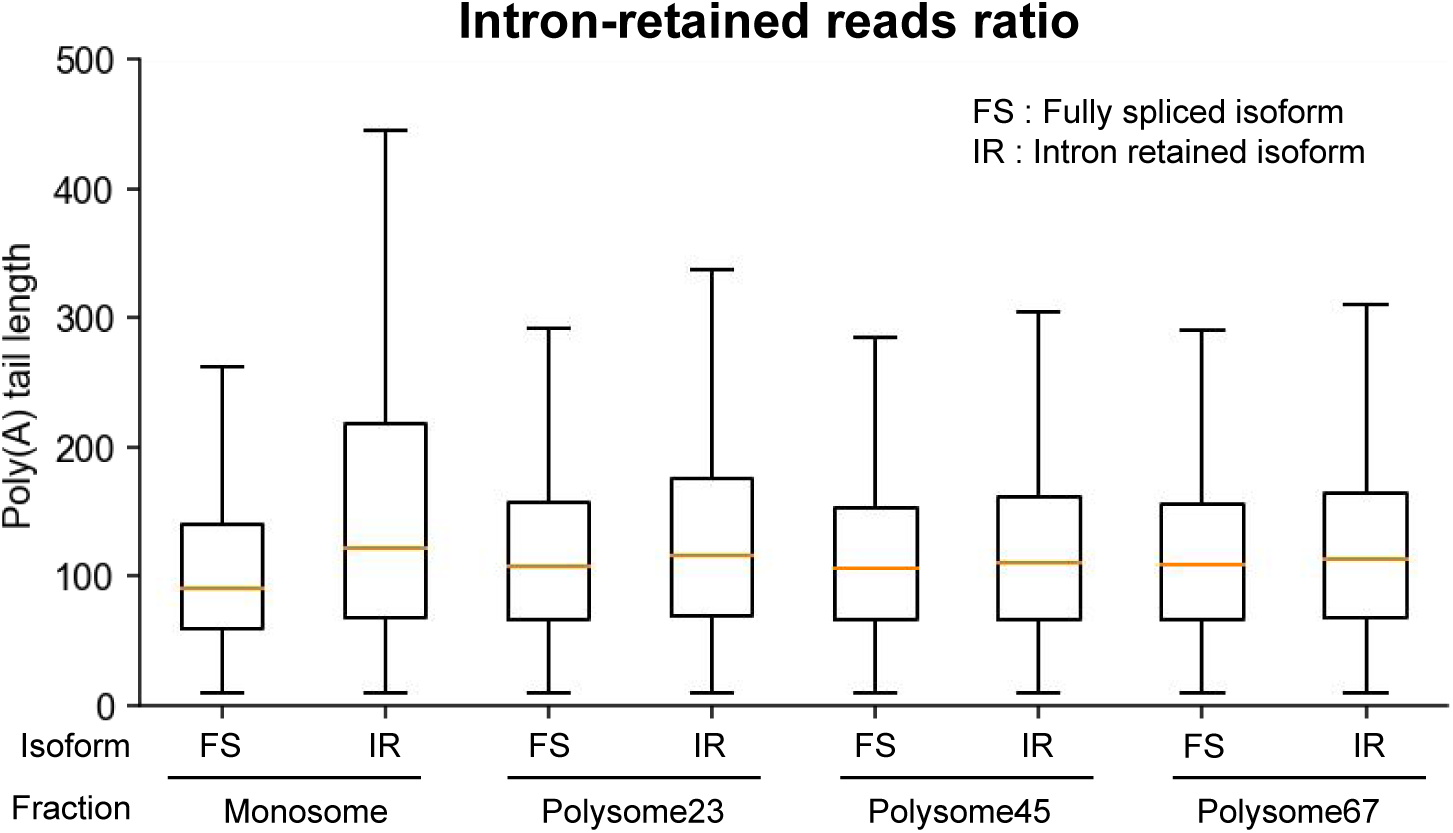
Comparison of poly(A) length in fully-spliced reads and Intron-retained reads in monosome, polysome23, polysome45, polysome67.

**Table S1.** Sequencing summary of SPARP-seq libraries.

| Libraries |  |  | Total reads | Aligned ratio | Aligned reads | mRNA reads | poly(A) reads |
| --- | --- | --- | --- | --- | --- | --- | --- |
| Nanopore | Monosome | rep1 | 17,240,321 | 93.48% | 17,309,388 | 6,893,286 | 6,141,610 |
|  |  | rep2 | 8,142,598 | 94.71% | 7,695,980 | 2,221,546 | 1,977,703 |
|  | Polysome23 | rep1 | 7,459,317 | 44.09% | 3,399,493 | 2,366,910 | 2,005,903 |
|  |  | rep2 | 12,801,194 | 95.28% | 12,170,378 | 6,851,740 | 4,844,011 |
|  | Polysome45 | rep1 | 8,332,407 | 45.57% | 3,917,570 | 3,176,018 | 2,510,120 |
|  |  | rep2 | 12,671,247 | 92.10% | 11,670,201 | 6,503,428 | 4,990,775 |
|  | Polysome67 | rep1 | 3,740,281 | 83.68% | 3,409,165 | 2,049,533 | 1,565,162 |
|  |  | rep2 | 16,350,537 | 91.82% | 15,013,211 | 6,750,711 | 5,240,989 |
| Pacbio | Monosome | rep1 | 252,444 | 99.57% | 251,358 | 175,208 | 128,953 |
|  | Polysome67 | rep1 | 272,415 | 99.31% | 270,535 | 215,404 | 149,440 |

**Table S2.** Sequencing summary of Ribo-seq libraries.

| Libraries |  |  | Total reads | Uniq mapped reads (24-37nt) |
| --- | --- | --- | --- | --- |
| Illumina | Monosome | rep1 | 13,137,888 | 1,505,112 |
|  |  | rep2 | 15,890,293 | 2,058,318 |
|  | Polysome | rep1 | 27,370,303 | 16,402,026 |
|  |  | rep2 | 24,942,196 | 14,801,992 |
|  | Total | rep1 | 32,071,825 | 21,317,866 |
|  |  | rep2 | 34,236,247 | 22,645,791 |

## References

1. Ingolia, N. T. Ribosome profiling: new views of translation, from single codons to genome scale. Nat. Rev. Genet. 15, 205–213 (2014).

2. Brar, G. A. & Weissman, J. S. Ribosome profiling reveals the what, when, where, and how of protein synthesis. Nat. Rev. Mol. Cell Biol. 16, 651–664 (2015).

3. Ingolia, N. T., Hussmann, J. A. & Weissman, J. S. Ribosome Profiling: Global Views of Translation. Cold Spring Harb. Perspect. Biol. 11, a032698 (2019).

4. Subtelny, A. O., Eichhorn, S. W., Chen, G. R., Sive, H. & Bartel, D. P. Poly(A)-tail profiling reveals an embryonic switch in translational control. Nature 508, 66–71 (2014).

5. Lima, S. A. et al. Short poly(A) tails are a conserved feature of highly expressed genes. Nat. Struct. Mol. Biol. 24, 1057–1063 (2017).

6. Eckmann, C. R., Rammelt, C. & Wahle, E. Control of poly(A) tail length. Wiley Interdiscip. Rev. RNA 2, 348–361 (2011).

7. Weill, L., Belloc, E., Bava, F.-A. & Méndez, R. Translational control by changes in poly(A) tail length: recycling mRNAs. Nat. Struct. Mol. Biol. 19, 577–585 (2012).

8. Lim, J., Lee, M., Son, A., Chang, H. & Kim, V. N. mTAIL-seq reveals dynamic poly(A) tail regulation in oocyte-to-embryo development. Genes Dev. 30, 1671–1682 (2016).

9. Nicholson, A. L. & Pasquinelli, A. E. Tales of Detailed Poly(A) Tails. Trends Cell Biol. 29, 191–200 (2019).

10. Passmore, L. A. & Coller, J. Roles of mRNA poly(A) tails in regulation of eukaryotic gene expression. Nat. Rev. Mol. Cell Biol. 23, 93–106 (2022).

11. Liu, J. & Lu, F. Beyond simple tails: poly(A) tail-mediated RNA epigenetic regulation. Trends Biochem. Sci. 49, 846–858 (2024).

12. Tudek, A. et al. Global view on the metabolism of RNA poly(A) tails in yeast Saccharomyces cerevisiae. Nat. Commun. 12, 4951 (2021).

13. Krawczyk, P. S. et al. Re-adenylation by TENT5A enhances efficacy of SARS-CoV-2 mRNA vaccines. Nature 641, 984–992 (2025).

14. Amrani, N., Ghosh, S., Mangus, D. A. & Jacobson, A. Translation factors promote the formation of two states of the closed-loop mRNP. Nature 453, 1276–1280 (2008).

15. Anger, A. M. et al. Structures of the human and Drosophila 80S ribosome. Nature 497, 80–85 (2013).

16. Sonenberg, N. & Hinnebusch, A. G. Regulation of Translation Initiation in Eukaryotes: Mechanisms and Biological Targets. Cell 136, 731–745 (2009).

17. Jackson, R. J., Hellen, C. U. T. & Pestova, T. V. The mechanism of eukaryotic translation initiation and principles of its regulation. Nat. Rev. Mol. Cell Biol. 11, 113–127 (2010).

18. Xiang, K., Ly, J. & Bartel, D. P. Control of poly(A)-tail length and translation in vertebrate oocytes and early embryos. Dev. Cell 59, 1058–1074.e11 (2024).

19. Xiong, Z. et al. Ultrasensitive Ribo-seq reveals translational landscapes during mammalian oocyte-to-embryo transition and pre-implantation development. Nat. Cell Biol. 24, 968–980 (2022).

20. Arava, Y. et al. Genome-wide analysis of mRNA translation profiles in Saccharomyces cerevisiae. Proc. Natl. Acad. Sci. U. S. A. 100, 3889–3894 (2003).

21. Vogel, C. & Marcotte, E. M. Insights into the regulation of protein abundance from proteomic and transcriptomic analyses. Nat. Rev. Genet. 13, 227–232 (2012).

22. Brito Querido, J., Irene Díaz-López & Ramakrishnan, V. The molecular basis of translation initiation and its regulation in eukaryotes. Nat. Rev. Mol. Cell Biol. 25, 168–186 (2024).

23. Heyer, E. E. & Moore, M. J. Redefining the Translational Status of 80S Monosomes. Cell 164, 757–769 (2016).

24. Biever, A. et al. Monosomes actively translate synaptic mRNAs in neuronal processes. Science 367, eaay4991 (2020).

25. Ingolia, N. T., Ghaemmaghami, S., Newman, J. R. S. & Weissman, J. S. Genome-wide analysis in vivo of translation with nucleotide resolution using ribosome profiling. Science 324, 218–223 (2009).

26. King, H. A. & Gerber, A. P. Translatome profiling: methods for genome-scale analysis of mRNA translation. Brief. Funct. Genomics 15, 22–31 (2016).

27. Ritter, A. J., Draper, J. M., Vollmers, C. & Sanford, J. R. Long-read subcellular fractionation and sequencing reveals the translational fate of full-length mRNA isoforms during neuronal differentiation. Genome Res. 34, 2000–2011 (2024).

28. Jagannatha, P. et al. Long-read Ribo-STAMP simultaneously measures transcription and translation with isoform resolution. Genome Res. 34, 2012–2024 (2024).

29. Ogami, K. et al. mTOR- and LARP1-dependent regulation of TOP mRNA poly(A) tail and ribosome loading. Cell Rep. 41, 111548 (2022).

30. Huang, S., Wylder, A. C. & Pan, T. Simultaneous nanopore profiling of mRNA m6A and pseudouridine reveals translation coordination. Nat. Biotechnol. 42, 1831–1835 (2024).

31. Viscardi, M. J. & Arribere, J. A. Poly(a) selection introduces bias and undue noise in direct RNA-sequencing. BMC Genomics 23, 530 (2022).

32. Legnini, I., Alles, J., Karaiskos, N., Ayoub, S. & Rajewsky, N. FLAM-seq: full-length mRNA sequencing reveals principles of poly(A) tail length control. Nat. Methods 16, 879–886 (2019).

33. Liu, Y., Nie, H., Liu, H. & Lu, F. Poly(A) inclusive RNA isoform sequencing (PAIso−seq) reveals wide-spread non-adenosine residues within RNA poly(A) tails. Nat. Commun. 10, 5292 (2019).

34. Begik, O. et al. Nano3P-seq: transcriptome-wide analysis of gene expression and tail dynamics using end-capture nanopore cDNA sequencing. Nat. Methods 20, 75–85 (2023).

35. Grandi, C. et al. Decoupled degradation and translation enables noise modulation by poly(A) tails. Cell Syst. 15, 526–543.e7 (2024).

36. Biziaev, N. et al. The impact of mRNA poly(A) tail length on eukaryotic translation stages. Nucleic Acids Res. 52, 7792–7808 (2024).

37. Weatheritt, R. J., Sterne-Weiler, T. & Blencowe, B. J. The ribosome-engaged landscape of alternative splicing. Nat. Struct. Mol. Biol. 23, 1117–1123 (2016).

38. Blencowe, B. J. The Relationship between Alternative Splicing and Proteomic Complexity. Trends Biochem. Sci. 42, 407–408 (2017).

39. Floor, S. N. & Doudna, J. A. Tunable protein synthesis by transcript isoforms in human cells. eLife 5, e10921 (2016).

40. Reixachs-Solé, M., Ruiz-Orera, J., Albà, M. M. & Eyras, E. Ribosome profiling at isoform level reveals evolutionary conserved impacts of differential splicing on the proteome. Nat. Commun. 11, 1768 (2020).

41. Cui, H., Hu, H., Zeng, J. & Chen, T. DeepShape: estimating isoform-level ribosome abundance and distribution with Ribo-seq data. BMC Bioinformatics 20, 678 (2019).

42. Thoreen, C. C. et al. A unifying model for mTORC1-mediated regulation of mRNA translation. Nature 485, 109–113 (2012).

43. Hu, W., Sweet, T. J., Chamnongpol, S., Baker, K. E. & Coller, J. Co-translational mRNA decay in Saccharomyces cerevisiae. Nature 461, 225–229 (2009).

44. Pelechano, V., Wei, W. & Steinmetz, L. M. Widespread co-translational RNA decay reveals ribosome dynamics. Cell 161, 1400–1412 (2015).

45. Webster, M. W. et al. mRNA Deadenylation Is Coupled to Translation Rates by the Differential Activities of Ccr4-Not Nucleases. Mol. Cell 70, 1089–1100.e8 (2018).

46. Monaghan, L., Longman, D. & Cáceres, J. F. Translation-coupled mRNA quality control mechanisms. EMBO J. 42, e114378 (2023).

47. Boothby, T. C., Zipper, R. S., van der Weele, C. M. & Wolniak, S. M. Removal of Retained Introns Regulates Translation in the Rapidly Developing Gametophyte of *Marsilea vestita*. Dev. Cell 24, 517–529 (2013).

48. Wong, J. J.-L. et al. Orchestrated Intron Retention Regulates Normal Granulocyte Differentiation. Cell 154, 583–595 (2013).

49. Ullrich, S. & Guigó, R. Dynamic changes in intron retention are tightly associated with regulation of splicing factors and proliferative activity during B-cell development. Nucleic Acids Res. 48, 1327–1340 (2020).

50. Chen, W., et al. Systematic characterization of the composition and dynamics of processing body-associated mRNAs. Preprint at 10.21203/rs.3.rs-5354706/v1 (2024).

51. Long, Y., Jia, J., Mo, W., Jin, X. & Zhai, J. FLEP-seq: simultaneous detection of RNA polymerase II position, splicing status, polyadenylation site and poly(A) tail length at genome-wide scale by single-molecule nascent RNA sequencing. Nat. Protoc. 16, 4355–4381 (2021).

52. Jia, J. et al. An atlas of plant full-length RNA reveals tissue-specific and monocots–dicots conserved regulation of poly(A) tail length. Nat. Plants 8, 1118–1126 (2022).

53. Iosub, I. A., Wilkins, O. G. & Ule, J. Riboseq-flow: A streamlined, reliable pipeline for ribosome profiling data analysis and quality control. Wellcome Open Res. 9, 179 (2024).

